# Interactive exploration of biobank-scale ancestral recombination graphs with Lorax

**DOI:** 10.64898/2026.02.19.706861

**Authors:** Pratik Katte, Russell Corbett-Detig

## Abstract

Ancestral Recombination Graphs (ARGs) underpin analyses of natural selection, disease association, and population history but remain difficult to explore at biobank scale. We introduce Lorax, a GPU-accelerated, web-native platform for real-time visualization of population-scale ARGs. Lorax integrates genomic position, coalescent time, local genealogy, and metadata, enabling interactive exploration of ancestry and variant inheritance in biobank-scale datasets.

## Background

Ancestral Recombination Graphs (ARGs) provide a complete description of ancestry within populations, encoding the full history of coalescence and recombination underlying observed genetic variation. As such, ARGs are increasingly used for inference of major evolutionary processes, including natural selection [1], demography [2], and admixture [3], among others [4-6]. Existing visualization tools, including the draw functionality in tskit [7], tskit-arg-visualizer [8], and ARGscape [9], lag behind ARG inference methods and are limited to small sample sizes and genomic regions. As a result, ARG visualization for biobank-scale datasets remains a major unsolved challenge that is limiting progress in biomedical and evolutionary genomics.

## Results and Discussion

To overcome the interactability and scalability bottleneck in ARGs visualization, we developed Lorax, a GPU-accelerated, web-native platform for interactive exploration of large-scale Ancestral Recombination Graphs. While tree sequences provide an extremely compact encoding of ARGs, Lorax’s central contribution is on-demand decoding and streaming of local genealogies for GPU rendering, allowing users to move seamlessly across genomic position, coalescent time, and local tree topology, while directly integrating sample metadata and variant annotations into the ancestry structure. This unified view allows users to identify population-specific genealogical patterns and trace variant inheritance through local genealogies, and scales to biobank-scale datasets, enabling interactive exploration of ARGs comprising millions of samples

Lorax integrates sample- and node-level metadata directly into the ARG visualization, enabling users to search, color, filter, and subset lineages by population labels, phenotypes, or other annotations. This makes population-specific coalescent patterns immediately visible, such as shallow, recent ancestry within a focal group contrasted with deeper shared ancestry across populations. Lorax also supports mutation-aware visualization, allowing the inheritance of specific variants to be traced through local genealogies, and includes genome-browser–compatible tracks for direct navigation from functional annotations to ancestry structure. Lorax additionally enables comparison of consecutive local topologies by highlighting where genealogical structure changes along the genome.

We demonstrate Lorax’s utility with two examples. First, we visualize human ancestry surrounding the well-studied lactase persistence locus near the LCT gene, a canonical case of recent strong positive selection in a subset of human populations. At this locus, Lorax reveals a cluster of predominantly European lineages that coalesce rapidly on the branch carrying the lactase persistence mutation rs4988235, consistent with a recent selective sweep associated with the adoption of dairy farming [10].

Second, we explore changing evolutionary relationships along the genomes of *Heliconius* butterflies, where a large chromosomal inversion is thought to have introgressed between species [11]. Lorax reveals a localized shift in genealogical structure within the inverted region, such that *H. sara, H. telesiphe*, and *H. demeter* which share the inversion, are more closely related within this genomic interval than elsewhere in the genome (Fig. 1b). Therefore, our tool can reveal deep-time evolutionary phenomena in addition to more recent population-level ancestral processes. More generally, these examples demonstrate Lorax’s ability to visualize both dense local genealogies and large-scale topological variation along the genome.

**Fig 1.**
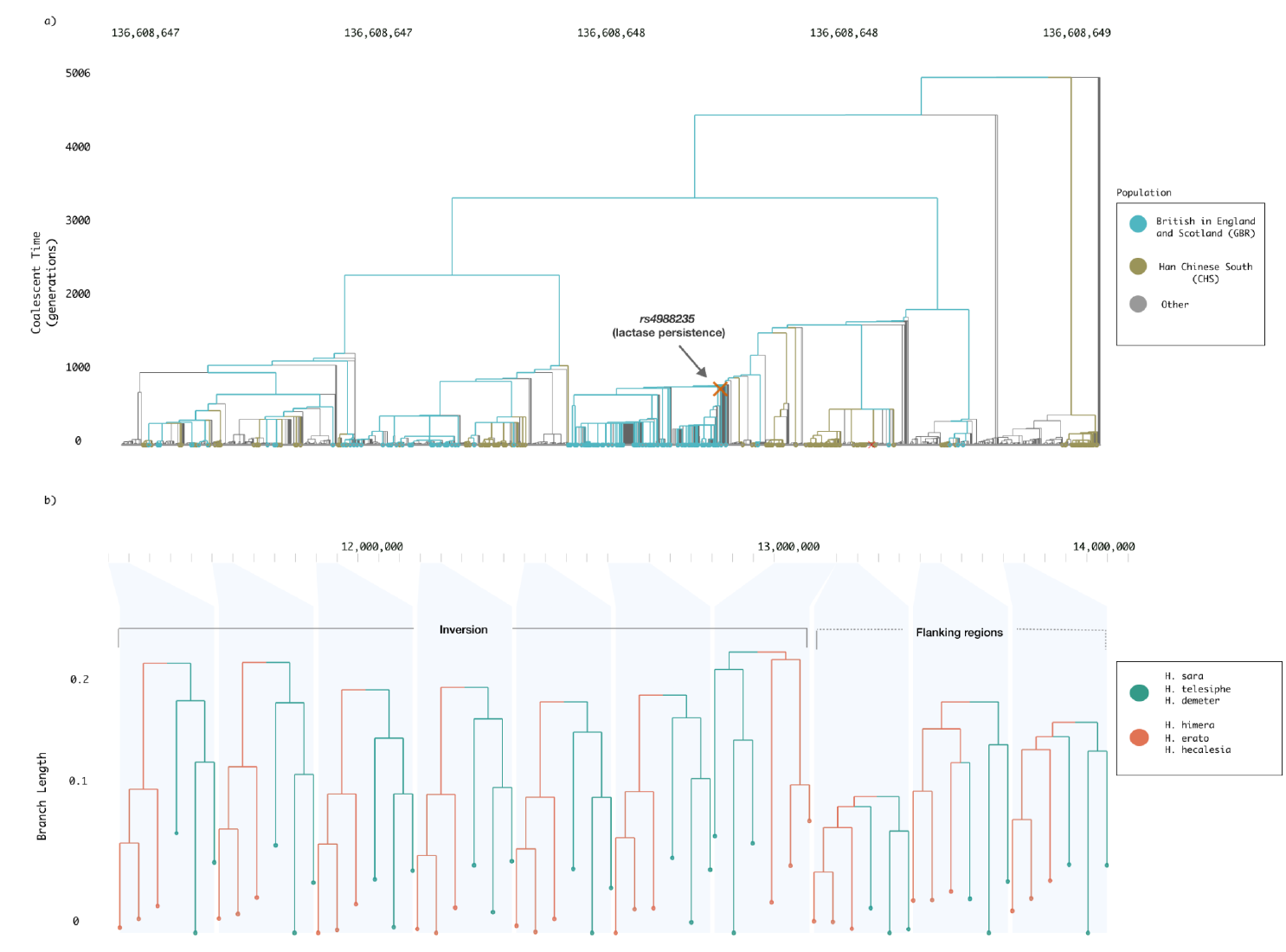
Interactive visualization of selection and introgression using Lorax. a) Genomic context of the lactase persistence locus. The regulatory variant rs4988235 lies within an intron of MCM6 and influences expression of the nearby LCT gene (link). b) Visualization of Heliconius chromosome 2 showing a localized shift in genealogical structure within a chromosomal inversion, where inversion-carrying samples are more closely related than in flanking genomic regions (link).

To demonstrate scalability for vast genomic datasets, we applied Lorax to the sc2ts[12] SARS-CoV-2 tree sequence dataset comprising ∼2.4 million viral sequences, one of the largest ARGs inferred to date, achieving real-time interactive rendering with mutation overlays and synchronized multi-view navigation (Supplementary Fig. 1). To systematically evaluate performance, we benchmarked Lorax on msprime-simulated [13] ARGs across a 50 Mb genomic region, varying diploid sample size from 200,000 to 1,000,000 and effective population size from 5,000 to 50,000, yielding tree sequences of ∼120,000 to 1.4 million local trees. Across all configurations, Lorax rendered initial trees within seconds and maintained memory use within practical bounds, confirming interactive performance at chromosome scale across the full range of biobank-relevant sample sizes (Supplementary Table 1).

## Conclusions

In conclusion, Lorax is a GPU-accelerated, web-based platform designed for interactive exploration of biobank-scale Ancestral Recombination Graphs. By enabling coordinated navigation across genomic position, time, and local genealogy with integrated metadata, Lorax makes complex ancestry structures directly interpretable. Lorax is open-source and freely available as both a pip package and a hosted web platform at https://lorax.ucsc.edu/.

## Methods

### Input Tree sequence and CSV file Format

Lorax supports tree sequence inputs in.trees format, its compressed.tsz variant, as well as CSV-based representations of ancestral recombination graphs and local genealogies. For.trees and.tsz files, Lorax uses the tskit Python library to extract sample and population metadata and access nodes and edges tables from tree sequence data structure enabling real-time reconstruction of local genealogies across genomic intervals. CSV inputs encode local genealogies on a per recombination interval basis, with each row specifying a genomic interval, a Newick-formatted tree, and a scalar summary of tree depth. These fields define the minimal structure required for rendering, while additional columns are treated as user-defined metadata.

### Implementation

Lorax uses a modular client–server architecture. A React-based frontend provides reusable components and custom GPU-accelerated deck.gl layers for visualizing local genealogies. A Python FastAPI backend processes uploaded files, manages persistent sessions, and supports low-latency interaction via Socket.IO. Computationally intensive tasks, including tree traversal and metadata queries are performed on the server side and streamed incrementally to the client using compact Apache Arrow IPC format, enabling real-time interactive rendering.

### Coordinated multi-view visualization

The Lorax interface uses multiple synchronized views that are rendered within a single canvas. A primary ARG view displays local genealogies, while aligned context views show genomic position, recombination interval markers along the horizontal axis, and coalescent time along the vertical axis. Interaction is coordinated by treating the main ARG view as the source of navigation state and further propagated to dependent views. This shared view state is mapped to explicit genomic coordinates that drive data retrieval and computation, ensuring that all visual elements remain consistent with the currently selected genomic region.

### GPU-accelerated rendering and metadata integration

The shared view state coordinates both data retrieval and rendering. As users navigate the genome, only local trees within the visible genomic window are requested from the backend, which streams compact binary payloads encoding traversal order, node relationships, and temporal bounds. On the client, a custom deck.gl layer renders precomputed branch and tip geometries as optimized WebGL primitives, operating directly on typed arrays to minimize per-frame overhead. Metadata is delivered through the same streaming channel for cohort-scale attributes. Together, these genealogical data and metadata streams are rendered through a single workflow enabling interactive filtering, coloring, search, and lineage tracing while minimizing network and rendering overhead at the scale of million samples.

## Declarations

### Competing Interests

RBC is a paid consultant for International Responder Systems.

### Ethics approval and consent to participate

NA

### Consent for publication

N/A

## Acknowledgements

The authors gratefully acknowledge helpful feedback from the Corbett-Detig lab and from members of the tskit development community. We also thank Jeff Ross-Ibarra for helpful feedback on preliminary versions of Lorax.

## Funding

This work was supported by National Institute of Health Grant number R35GM128932 to RBC.

## Code Availability

Lorax is an open-source tool available both as a [pip package](https://pypi.org/project/lorax-arg/) and as a hosted web platform at http://lorax.ucsc.edu/. The source code and full documentation are available on [GitHub](https://github.com/pratikkatte/lorax/). The scripts used to preprocess the Heliconius dataset and to generate the Supplementary performance table are included in the scripts/ directory of the repository.

## Data Availability

The 1000 Genomes tree sequence used to visualize the lactase persistence locus is publicly available from Zenodo (https://zenodo.org/records/3051855). Genomic data for Heliconius butterflies are available from Dryad (https://doi.org/10.5061/dryad.b7bj832), as described in Edelman et al. [11]. No new experimental data were generated in this study. Processed files used in the demonstrations are available through the Lorax web platform at https://lorax.ucsc.edu.

## Authors’ contributions

P.K. developed the Lorax tool under the supervision of R.C. P.K and R.C co-wrote the manuscript.

## Notes

https://lorax.ucsc.edu/

https://github.com/pratikkatte/lorax/

https://pypi.org/project/lorax-arg/

