## Supplemental Material for "Interactive exploration of biobank-scale ancestral recombination graphs with Lorax"

**Supplementary Fig 1 | Interactive rendering of a large SARS-CoV-2 tree-sequence dataset in Lorax.**

Lorax view of local genealogies from the sc2ts dataset, demonstrating real-time rendering and navigation at large scale (~2.4 million sequences). This panel illustrates dense local-tree structure, mutation overlays, and synchronized coordinate-linked visualization under high data volume.

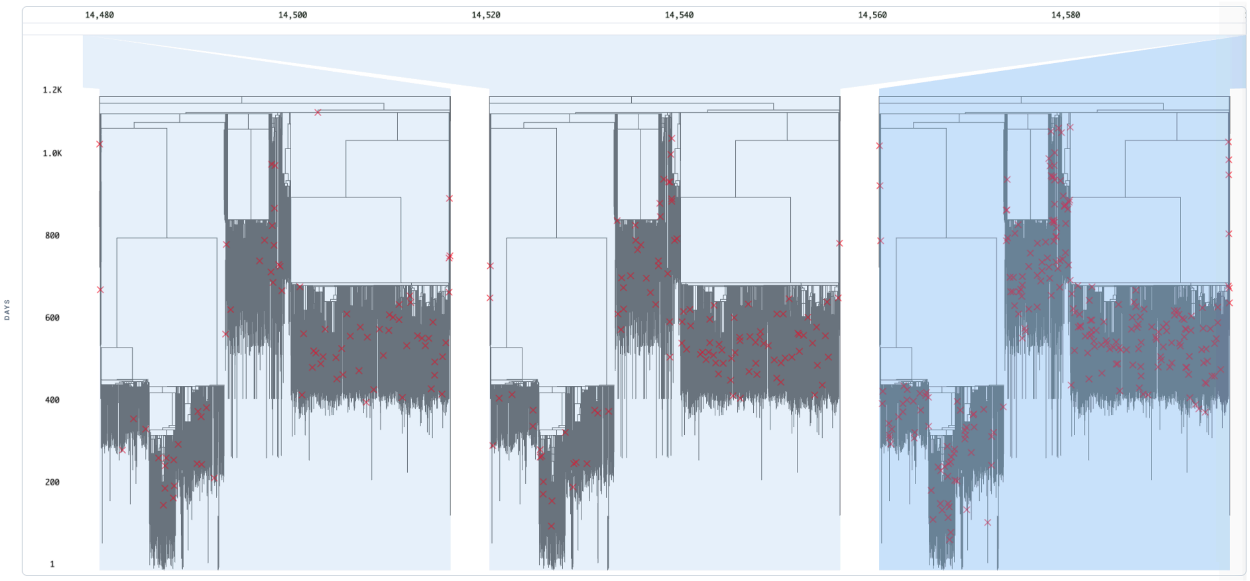


**Supplementary Table 1 | Lorax performance across diploid sample size and effective population size ($N_e$)**

| Diploid Individuals | $N_e$ | File size (MB) | Local trees | Mutation | Load time (s) | First layout (s) |
|---|---|---|---|---|---|---|
| 200,000 | 5,000 | 76M | 127,802 | 168,611 | 1.52 | 2.77 |
| | 50,000 | 311M | 1,263,843 | 1,683,806 | 3.75 | 3.11 |
| 600,000 | 5,000 | 126M | 138,754 | 182,560 | 4.00 | 9.73 |
| | 50,000 | 432M | 1,372,235 | 1,822,947 | 6.79 | 10.53 |
| 1,000,000 | 5,000 | 275M | 144,110 | 188,937 | 6.56 | 18.42 |
| | 50,000 | 541M | 1,424,671 | 1,890,836 | 9.41 | 16.79 |
